# Substrate heterogeneity promotes cancer cell dissemination through interface roughening

**DOI:** 10.1101/2025.05.20.655037

**Authors:** Zuzana Dunajova, Saren Tasciyan, Juraj Majek, Jack Merrin, Erik Sahai, Michael Sixt, Edouard Hannezo

## Abstract

While tumor malignancy has been extensively studied under the prism of genetic and epigenetic heterogeneity, tumor cell states also critically depend on reciprocal interactions with the microenvironment. This raises the hitherto untested possibility that heterogeneity of the untransformed tumor stroma can actively fuel malignant progression. As biological heterogeneity is inherently difficult to control, we adopted a reductionist approach and let tumor cells invade micro-engineered environments harboring obstacles with precision-controlled geometry. We find that not only the presence of obstacles, but more surprisingly their spatial disorder, causes a drastic shift from a collective to a single-cell mode of invasion – comparable in strength to cadherin loss. Combining live-imaging and perturbation experiments with minimal biophysical modeling, we demonstrate that cell detachments result both from local geometrical constraints and a global integration of spatial disorder over time. We show that different types of microenvironments map onto different universality classes of invasion dynamics - homogeneous substrates follow Kardar–Parisi–Zhang (KPZ) scaling, while disordered ones exhibit exponents consistent with KPZ with quenched disorder (KPZq). Our findings highlight generic physical principles for how the mode of cancer cell invasion depends on environmental heterogeneity, with potential implications to understand tumor evolution *in vivo*.

## Introduction

Cellular heterogeneity is at the core of tumor evolution. The canonical view suggests that new genetic variants produce new cancer cell phenotypes that invade extra-tumoral micromilieus. When such variants experience favorable conditions, they clonally expand, amplifying genetic heterogeneity in an iterative process that ultimately leads to a fatal state of systemic dissemination^1–3^. While heritable genetic variations have long been recognized to play a crucial role, phenotypic variation can also arise from epigenetic regulation or transient signaling states^4,5^. Importantly, variation and selection of cancer cells are not autonomous to the tumor, but occur through reciprocal crosstalk with the untransformed microenvironment ^6–8^. Remarkably, in some tumor models, invasiveness is not an intrinsic property of tumor cells but rather a transient response of rare cancer cells to signals released from a benign stroma^9–11^. This raises the possibility that heterogeneous extra-tumoral microenvironments may act as drivers of malignancy^12^, although this remains largely untested due to the experimental difficulty of controlling substrate heterogeneity.

In recent years, reductionist approaches have gained increasing attention, showing that cell behaviors can emerge from physical principles without requiring genetic or epigenetic changes. For instance, bottom-up reconstitution of microenvironments has been successfully used to model cellular responses^13^ and cancer dissemination in response to specific extracellular matrix (ECM) topographies^14,15^. Cellular collectives can also undergo fluid-solid transitions as a function of cell-cell and cell-ECM adhesion, cell density or active migratory forces^16–22^, with fluidized states favoring invasion^23–27^. Such fluidization can create large mechanical stresses on nuclei, triggering mechano-adaptative responses that feed back to further promote malignancy^28^. In parallel, the interfaces of spreading tumor colonies display fractal-like features, that have been proposed to follow generic models for growing interfaces, such as the Kardar-Parisi-Zhang (KPZ) equation^29,30^ or the Molecular Beam Epitaxy (MBE) equation^31^, validated in diverse non-living systems^32,33^. However, prior studies have almost exclusively examined migration in homogeneous or ordered environments, leaving the impact of extracellular heterogeneity unexplored.

Here, we combine reductionist microengineering pillar arrays, live cell imaging and minimal particle-based simulations to dissect how environmental disorder shapes collective invasion. We chose to specifically manipulate environmental geometry, while leaving all other aspects, like cell-cell and cell-matrix adhesions, constant. This minimal change demonstrated complex impacts on interfacial growth and tumor dispersion. In particular, we found that spatial disorder robustly enhances single-cell detachments from the invasive front, a phenomenon that emerges both from local features of substrate heterogeneity, as well as a much more global ef-fect of temporal integration of the heterogeneity causing the roughening of the tumor front. Indeed, substrate heterogeneity leads to a transition to a different universality class of invasion dynamics, which we characterize experimentally and theoretically. This led us to derive a quantitative description of the determinants of cellular dispersion, which we tested by experiments in different environments with structured geometries. Our results link microenvironmental heterogeneity to invasion dynamics through universal physical principles, providing a mechanistic basis for how disorder promotes cellular dispersion and, potentially, malignant progression.

## Results

### Environmental mechanics and heterogeneity are sufficient to drive dispersion *in vitro*

As a cellular model for tumor invasion, we employed A431 human carcinoma cells^34^. This cell line is derived from a primary tumor of epithelial origin and forms stable cell-cell adhesions due to the expression of E- and P-Cadherin^35^. On fibronectin-coated 2D surfaces, A431 cells spread radially as a cohesive front, without cellular detachments (Supplementary Fig. 1a). To test invasion into geometrically precisely controlled environments, we engineered microfluidic devices where a centrally applied pool of cells spread into a fibronectin-coated confined space of 9μm height. The two confining surfaces were intersected with arbitrarily positionable 9μm diameter pillars, creating a 3D chamber that forces cells to invade into the “pillar forest” as a monolayer. This 9μm pillar spacing was chosen both so that cells could pass through by only moderately deforming their ∼12-18μm diameter nuclei, consistent with dimensions of 3D ECM pores *in vivo* ^36,37^. The setup allowed free medium exchange and long-term life imaging (>1week) using genetically encoded fluorescent reporters for cytoplasm (mCherry) and nuclei (GFP).

Each microfluidic chip was divided into four sectors, allowing parallel testing of different pillar patterns under identical conditions. The ordered patterns consisted of grid-like arrays of obstacles with constant spacing (Fig. 1a), whereas in disordered patterns the average spacing was preserved, but individual pillar positions randomly shifted according to a Laplace distribution (Fig. 1b). We define the level of spatial disorder by the average displacement of pillars relative to an ordered array (see Methods). Tracking invasion over 8 days showed that cells invaded ordered patterns with occasional detachments. Strikingly, in disordered patterns cells advanced with enhanced single cell detachments at the invasion front (Mann–Whitney U test, p = 0.0075, Fig. 1c,e, Supplementary Movie 1). This was in stark contrast to experiments on obstacle-free 2D surfaces, where A431 cells remained cohesive with no detachments (Fig. 1d, Supplementary Movie 2).

**Fig. 1:**
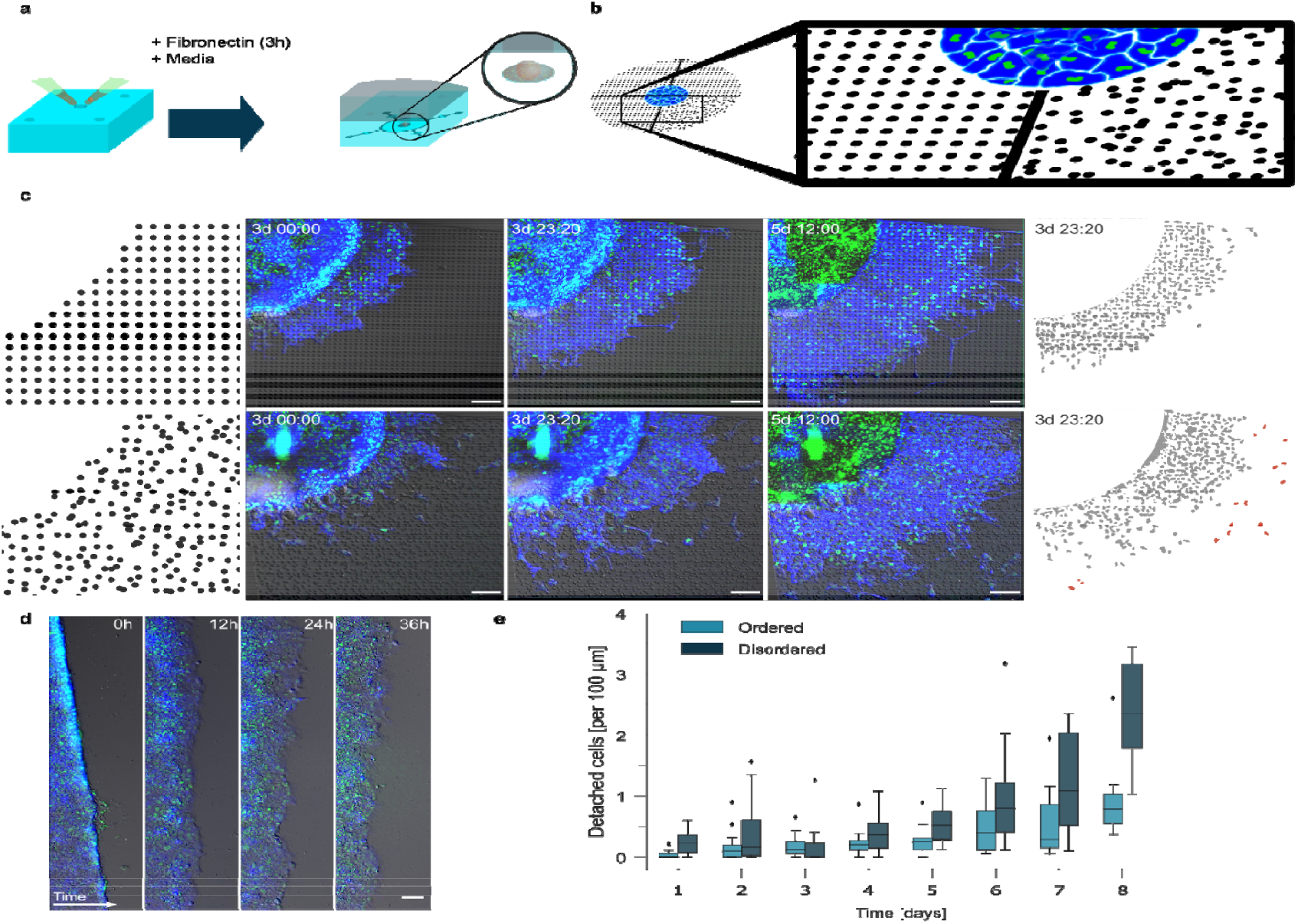
Substrate heterogeneity promotes tumor cell dispersal *in vitro*. **a**. Experimental setup of the microfluidic invasion assay. Devices are coated with fibronectin and filled with cell culture media before A431 cells are seeded at the center **b**. Illustration of the ordered and disordered pillar patterns used to modulate substrate geometry. Cells invade outward from the central seeding region. **c**. A431 cell invasion imaged with time-lapse fluorescence microscopy over the days 3, 4 and 5.5 from left to right. Cells express cytoplasmic mCherry (blue) and nuclear H2B-GFP (green). Right panel shows segmented nuclei at day 4. **d**. Scratch assay in homogeneous (pillar-free) environment, serving as a control for collective migration. **e**. Number of detached cells normalized by the radial interface length in the respective pillar patterns for A431 WT. Statistical comparison (Mann–Whitney U): Ordered vs. Disordered, p = 0.0075. Scale bars: 100 μm, box and whiskers plot display median, 1^st^ and 3^rd^ quartiles (hinges) and bars display 1.5 interquartile range from the hinges, n=16 experiments for ordered and n=18 per disordered environment, followed over 8 days. These data show that substrate disorder significantly increases the frequency of single-cell detachment events, despite identical average pore size and cell type, indicating a direct role for environmental geometry in driving dispersal.

To further asses the magnitude of the dispersion effect in disordered patterns, we compared wild-type (WT) A431 cells to cells that were genetically engineered to lack E- and P-cadherin (EPCad^-/-^). As expected, EPCad^-/-^ cells invaded primarily as single cells, irrespective of environmental geometry (Supplementary Fig. 1b-c, Supplementary Movie 3). Notably, WT cells in disordered environments showed detachment frequencies comparable to EPCad^-/-^ cells (Supplementary Fig. 1d). Together, these data demonstrate that substrate disorder alone can drive a drastic shift from collective to single cell invasion, with an effect comparable in magnitude to genetically inflicted loss of cell-cell adhesion.

Several mechanisms might explain why environmental heterogeneity promotes single cell detachments, as different geometrical features change in ordered vs. disordered patterns. Firstly, we reasoned that disordered pillar patterns contain regions of very narrow pore sizes, that mechanically restrict cell-cell contacts. Such constrictions could also compress the nucleus, potentially triggering an active cellular response that might propel the cell forward and cause detachment^38–43^. To test this ‘pore size’ hypothesis, we repeated the assay in ordered pillar patterns with smaller spacing (6μm), forcing nuclei to severely squeeze between obstacles. We found that cells nonetheless invaded collectively without increased detachment, arguing against pore size alone as the cause (Supplementary Fig. 1d).

Next, we hypothesized that gradients in pore size, present only in disordered environments, could promote detachments. In this “funnel” scenario, cells at the leading edge might pass through a narrow constriction into more open space, while the bulk of the tissue remains stalled. To test this, we engineered two ordered but graded pillar arrays, in which pore size varied only along the invasion axis; one gradual (sinusoidal design) and one abrupt (stepwise design) (see Supplementary Fig. 1e). Importantly, neither design triggered a significant increase in detachments (Mann–Whitney U test: p = 0,61 and p = 0.88, respectively; Supplementary Fig. 1f). Together, these data show that favorable local obstacle geometry alone is not sufficient to increase cellular detachments. This led us to explore other, more collective mechanisms, where repeated transitions through heterogeneous configurations accumulate to drive detachment in disordered environments.

### Minimal biophysical simulations recapitulate the effect of environmental heterogeneity

To further test the role of disorder, we turned to *in silico* simulations of invasion in complex environments. We deliberately adopted a minimal modeling approach, asking whether spatial disorder alone, in the absence of complex cell-intrinsic features, was sufficient to promote cell detachment. Cells were modeled as soft, adhesive, and polar self-propelled particles^44–47^, interacting repulsively with static pillars arranged in different geometries (Fig. 2a). To mirror our analysis of experimental data, detached cells were defined in the simulations as cells separated by distance 1.55 *L* from the bulk, i.e. when the cell-cell contact force becomes negligible (see Methods for details). Before quantitatively constraining the model parameters, we first performed a broad parameter screen. Despite the model’s simplicity, it robustly recapitulated key qualitative features of the experiments: detachments increased strongly with the presence of pillars, and even more so with pillar heterogeneity (Supplementary Fig. 2a). These features were robust to model variations, for example, adding transient cell-pillar adhesion did not change the phenomenology (Supplementary Fig. 2b).

**Fig. 2:**
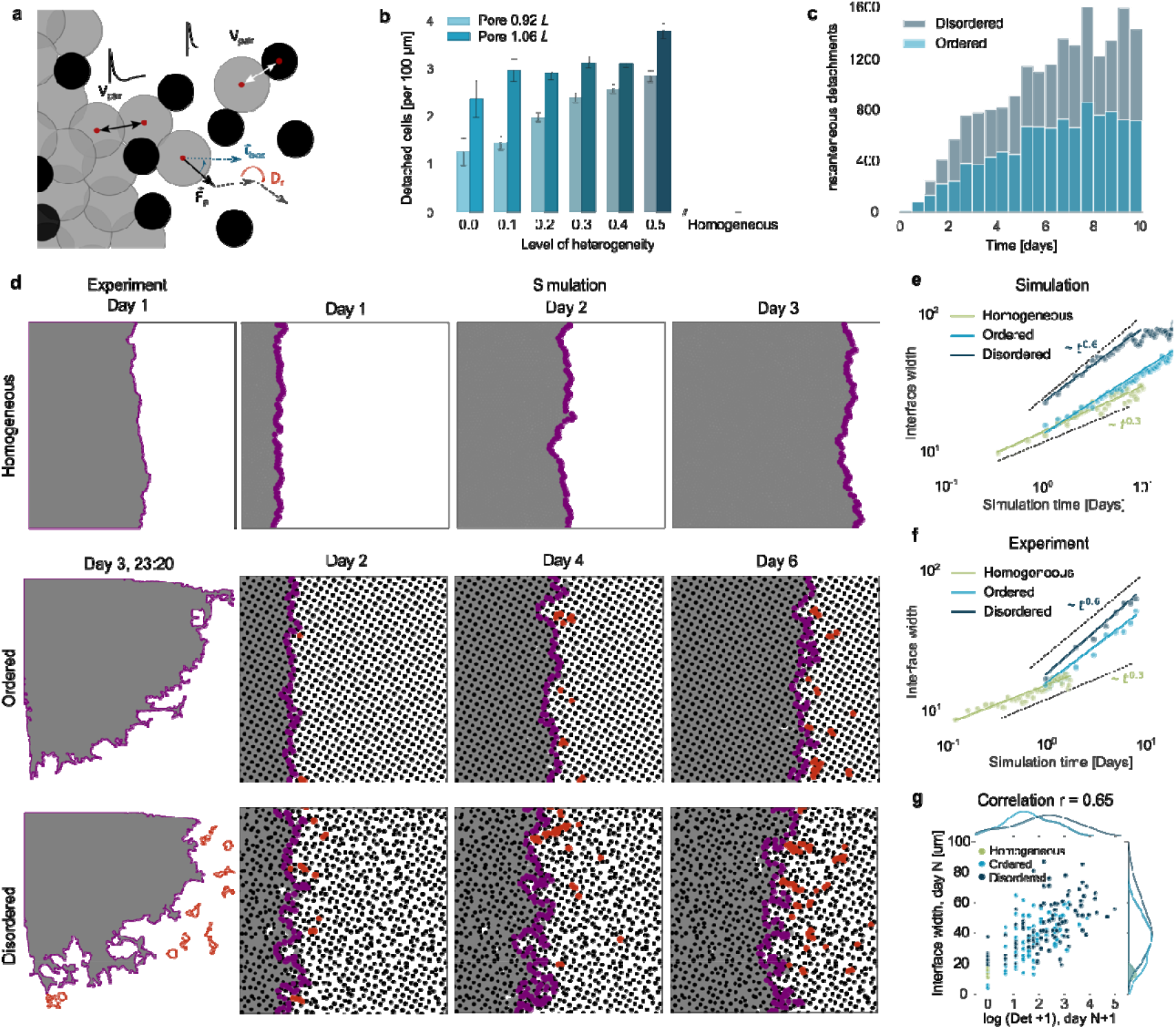
Minimal particle-based simulations reproduce heterogeneity driven detachments and reveal differences in dynamics of interfacial growth. **a**. Schematic of the simulation setup. Cells are modeled as soft, adhesive particles (gray) invading arrays with pillar obstacles (black). Relevant forces and interaction potentials are illustrated. **b**. Simulated number of detached cells (at day 6) as a function of environmental disorder. Disorder is controlled via the scale parameter of a Laplace distribution (see Supplementary Table 1). Bars show mean values; error bars indicate standard error. Increase of detachment frequency was approximated by ordinary least squares (OLS) regression model: 3.31 ± 0.26 (R^2^ = 0.98, p < 10^-4^) and 2.19 ± 0.51 (R^2^ = 0.82, p = 0.013) for pore sizes of 0.92□*L* and 1.06□*L*, respectively. **c**. Rate of detachment events increases over time in simulations, with steeper increase in disordered patterns. OLS regression slopes are 2.28 ± 0.19 (R^2^ = 0.89, p < 10^-4^), and 4.29 ± 0.32 (R^2^ = 0.91, p < 10^-4^) for ordered and disordered, respectively. **d**. Snapshots of simulated invasion fronts at different time points (magenta), alongside experimental cell interface segmentations (see Fig. 1c,d). Interface roughness increases over time and with environmental disorder. Detached cells are shown in red. **e**. Quantification of interfacial roughness over time in simulations. Power-law fits yield growth exponents of *β*_hom_= 0.33 ± 0.02, *β*_ord_= 0.46 ± 0.01, *β*_dis_ = 0.53 ± 0.02 (R^2^ > 0.92, p < 10^-4^), consistent with increasing roughening in disordered environments. Markers indicate means, shaded regions show standard error and solid lines the linear fits. **f**. Quantification of interfacial roughness in experiments yields growth exponents *β*_hom_= 0.25 ± 0.01, *β*_ord_ = 0.54 ± 0.04, *β*_dis_= 0.63 ± 0.04 (weighted least squares regression, R^2^ > 0.92, p < 10^-4^), showing a good agreement with the theoretical predictions. **g**. Across all experiments, interface roughness at day□N strongly correlates with the number of detachments observed on day□N+1 (n_ord_= 16; n_dis_=18, n_hom_= 6, Pearson r = 0.66, p < 10^-4^). Together, these results show that increasing substrate disorder leads to stronger interfacial roughening and elevated detachment rates, both in simulations and experiments.

We then constrained model parameters more precisely, to understand quantitatively what causes detachments in the simulations. As detailed in the Methods (see also Supplementary Table 1 for a summary of the parameter fitting), parameters governing single-cell migration (e.g. migration speed and bias, persistence time) were inferred from the measured behavior of single detached cells in pillar patterns. Cell-cell adhesion strength was constrained based on the lack of detachments in homogeneous environment and infrequent cell-cell rearrangements, consistent with a glass-like rheology (Supplementary Fig. 2c, Supplementary Movie 4). Simulations within this parameter regime confirmed that pillar heterogeneity consistently increased detachments (OLS regression, R^2^ = 0.98 and R^2^ = 0.82 for pore sizes 0.92 and 1.06 *L*, respectively – see Fig. 2b, Supplementary Movie 5). Furthermore, reducing adhesion strength not only increased detachments overall, but also diminished the additional effect of pillar heterogeneity (T-test: p=0.47; Supplementary Fig. 2d,e, Supplementary Movie 6), in good agreement with experimental results from EPCad^-/-^ cells (Supplementary Fig. 1d).

With the constrained parametrization, we next ran many repetitions of simulations with a specific disordered pattern and calculated the probability distribution of detachment as a function of position within the pillar pattern. This analysis, based on clustering instantaneous cell positions and tracking both transient and long-term detachments, revealed that detachment probability was most strikingly correlated with time, rather than a specific local position. Indeed, the frequency of detachments consistently increased (OLS regression: R^2^ = 0.9, p < 10^-4^) as the invasion progressed, with much steeper increase in disordered patterns (Fig. 2c).

### Substrate heterogeneity drives KPZq-type roughening of tumor invasion fronts

From a physical perspective, a classical result on the invasion dynamics of a fluid material such as a tissue is that its interface becomes progressively rougher over time, following a highly universal trend^29,48–51^, which depends only on a few key characteristics of the system. We hypothesized that cell detachments could be connected to these interface properties, with “time-integrated” heterogeneity causing cumulative surface roughening, as each pillar perturbs the advancing front.

To test this, we computed in simulations both the overall interface perimeter (which increased continuously during cell invasion, Supplementary Fig. 3a), as well as the surface roughness (i.e. deviations in interface height from its mean, Fig. 2d,e). Interestingly, the interface roughness always increased in time, but the growth exponent depended strongly on geometry. Power-law fits yielded *β*= 0.33 ± 0.02 in homogeneous environments – consistent with KPZ dynamics, a highly universal model of interfacial growth. Strikingly, disordered patterns showed markedly faster roughening with □ = 0.5-0.6. This is close to predictions for the KPZq universality class^52–55^, which describes interfaces in the presence of quenched noise (Fig. 2e). Given that pillars are static in our simulations, they can indeed be considered as quenched noise sources for the migrating interface.

Computing □ across a wide parameter space (Supplementary Table 2) confirmed that these exponents were robust despite the additional features absent from continuum KPZ models (e.g. adhesion and self-propulsion). To further test identification of the universality classes, we independently analyzed the height–height correlation function *C*(*r*)= ⟨[*h*(*x*+*r*) − *h*(*x*)]^2^⟩∼*r*^2*α*^ to extract the roughness exponent *α* ^56,57^. In homogeneous environments, *α* = 0.54 ± 0.03, again consistent with KPZ scaling (*α* ≈ 0.5, Supple-from the simulations at late times, mentary Fig. 3b). In disordered environments increased to 0.71 ± 0.01, which is in agreement with the KPZq universality class^57^. Finally, we asked whether the measured exponents jointly satisfy the Family–Vicsek scaling relation^48^, which predicts that data from different lateral system sizes collapse onto a universal curve when roughness and time are rescaled as *w*/*L*^*α*^ and *t*/*L*^*α*^ respectively. Using the independently extracted exponents (*α*≈ 0.55, *β* ≈ 0.33 in homogeneous and *α* ≈ 0.70, *β*≈ 0.55 in disordered environments), we observed a very good collapse across system sizes (Supplementary Fig. 3c). This stringent scaling test^58^ provides strong evidence that environmental disorder fundamentally alters the universality class of collective cell invasion.

**Fig. 3:**
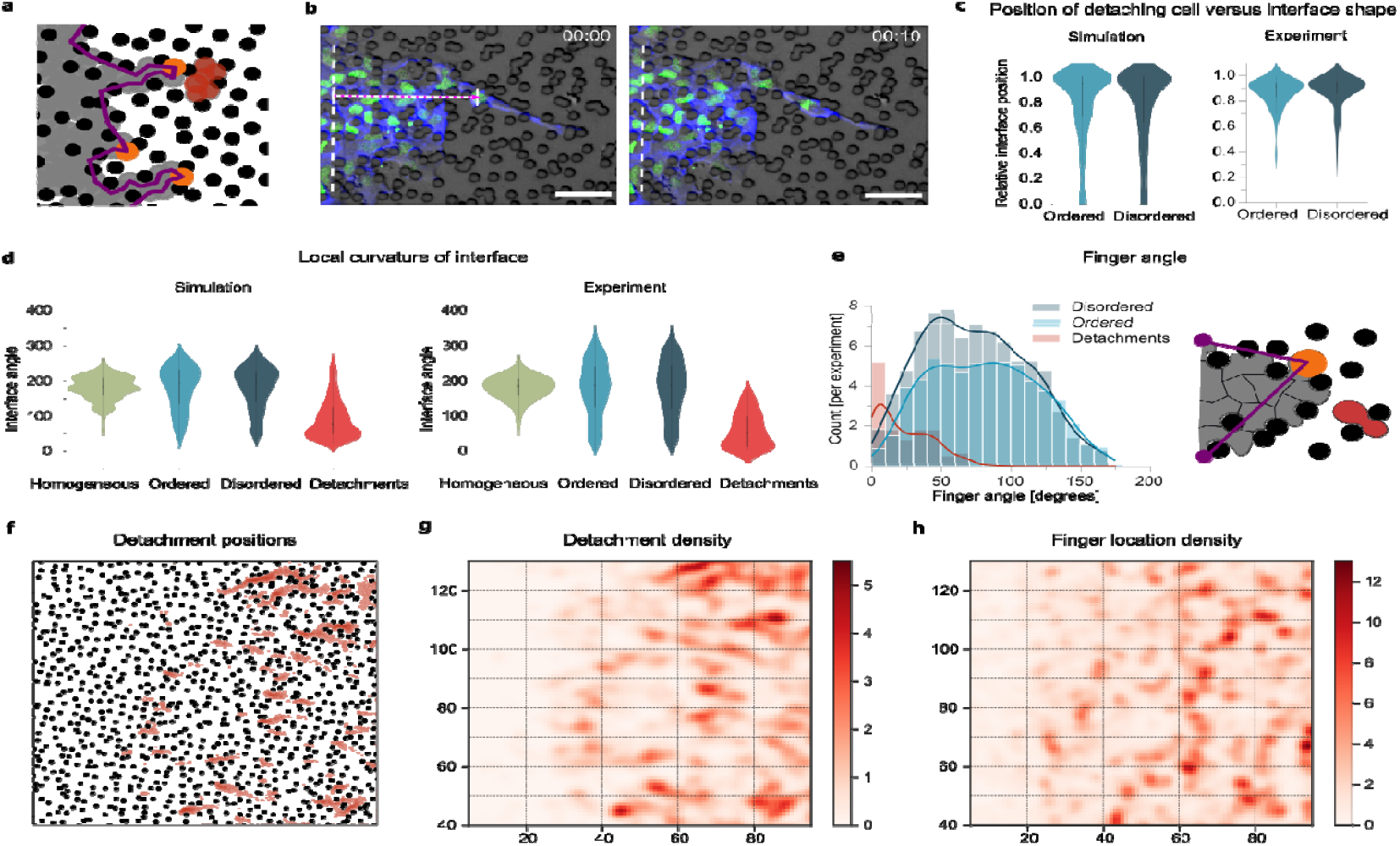
Local interface geometry predicts cell detachment across conditions. **a**. Interface shape (magenta) analyzed in simulations. Cells are shown as grey and pillar as black beads, tips of interface protrusions are highlighted in orange, detached cells in red. **b**. Schematic of how the position of detaching cells is measured relative to interface shape in both simulations and experiments. Detachment positions are normalized between the base (0) and tip (1) of the nearest protrusion. The last time frame prior to detachment is used. Cell bodies express cytoplasmic mCherry (blue) and nuclear H2B-GFP (green). Scale bar, 100µm. **c**. Distribution of detachment positions relative to the interface shape. Detachments predominantly occur near protrusion tips across all conditions. Middle line, upper and lower lines indicate the median, first and third quartiles, respectively (collection of n=100 detaching cells from n=7 experiments and n>700 from n=40 simulations for each condition). **d**. Local curvature distributions of the interface in homogeneous, ordered and disordered environments, compared with curvature at detachment points. Pillar patterns, especially disordered, show a higher frequency of large-curvature regions. Data from 6 (homogeneous), 16 (ordered), and 18 (disordered) experiments, and 40 simulations per condition. **e**. Aspect ratio of interface protrusions in experiments, defined by the angle between the tip and the adjacent local minima (“shoulders“). Disordered environments produce more elongated, narrow protrusions (Ordered vs. Disordered distribution, p < 10-4). Same dataset was used as in d). **f-h**. Spatial distribution of interfacial protrusions (h) and detachments (g) across n=10 simulations with identical disordered pillar pattern (f), confirming that the location of interface protrusions is highly predictive of detachments (Pearson r = 0.6, p < 10-4). These results show that local geometric features - especially sharp, convex interface protrusions - strongly predict detachment events in both experiments and simulations.

Strikingly, computing surface roughness and the growth exponent *β* in the experimental data revealed strong agreement with the simulations (*β* =0.25 ± 0.01 in homogeneous and *β* =0.63 ± 0.04 in disordered environments, Fig. 2f). In simulations of ordered pillar patterns, we found that the predicted growth exponent depended both on the average pore size and the orientation of the pattern relative to invasion direction (Supplementary Table 2, Supplementary Fig. 3d,e). Typically, ordered environments yielded intermediate exponents, with systematically lower interface width than disordered environments. This trend was confirmed experimentally: ordered patterns exhibited *β* =0.54 ± 0.04, and the average interface width was 25% lower compared to disordered environments (Mann-Whitney U test, p=0.0004).

Having rationalized the dynamics of interfacial growth, we next asked whether this could explain the different trends of temporal increase in detachments across conditions. To test this, we correlated interface roughness at day N with the number of detachments at day N+1 for each experimental replicate. We found a strong correlation (Pearson r = 0.66, p < 10^-4^) across all conditions, arguing that there might be a causal link between interfacial roughening and cell detachments (Fig. 2g), suggesting a direct mechanistic link between roughening and detachment. To test further this idea, we revisited geometries that did not produce detachments, such as ordered patterns with small pores and sinusoid/stepwise designs (Supplementary Fig. 3f, Supplementary Movie 7). Simulations run under the same parameter regime but with these pillar geometries reproduced the experimental outcome: both interfacial roughness and number of detachments remained lower than in disordered environments (Supplementary Fig. 3g). Together, these results indicate that substrate disorder promotes detachments by enhancing interface roughening, motivating us to next ask where along the interface detachments preferentially occur and which local geometrical features make them more likely.

### Quantitative local and global determinants of cellular detachments

To follow up on this finding, we asked how a global property such as interface roughness connects to the local occurrence of cell detachments. We hypothesized that rougher fronts generate interface regions with higher curvature and fewer cell contacts, which might facilitate detachments^59–61^. Indeed, analysis of individual detachment events in both simulation and experiment showed that the cells were most likely to detach from the tip of convex, “finger-like” protrusions (Fig. 3a-c). We next analyzed the local curvature of the interface immediately before detachment, which is inversely related to the number of neighbors (Fig. 3d). Most detachments occurred from interface positions above a threshold curvature value. By contrast, cells migrating in pillar-free environments never reached such high interfacial curvature values, quantitatively explaining the absence of detachment in this condition. Although the shape of curvature distributions of ordered and disordered patterns appear similar in experiments, likely due to the challenges of segmenting very rough interfaces (Supplementary Fig. S4a), we found that the disordered patterns yield 50 % more sharper protrusions than ordered ones (finger angle < 50°, p < 10^-4^), especially at late times (Fig. 3e, Supplementary Fig. 4b). Our theory was further supported by a strong spatial relationship between detachments and interfacial protrusions created by the heterogeneous environment (Pearson r = 0.6, p < 10^-4^; Fig. 3f-h, Supplementary Fig. 4c). Together, these findings support the idea that cell dispersion in pillar patterns is a cumulative phenomenon. Cell front invades with gradual increase of interfacial roughness, with geometry-dependent scaling exponents. The existence of a critical roughness which is permissive for detachments explains that detachments do not occur at early times in the pillar patterns and never in the pillars-free conditions (within our experimental window).

To test this mechanism further via independent experiments, we designed new pillar geometries consisting of sinusoidal patterns oriented perpendicular to the invasion axis. By making the sinusoidal modulation relatively mild, we ensured invasion across the entire pillar pattern while systematically controlling curvature modulation (Fig. 4a, Supplementary Movie 8). As predicted by simulations, this geometry promoted formation of periodic finger-like protrusions at re-gions of maximal pore width (Fig. 4b), and to a higher detachment probability when the frequency of sinusoidal modulation increased, producing sharper fingers (Supplementary Fig. 4d). In both simulations and experiments, the detachment probability scaled with the local interface shape, confirming that curvature modulation alone is sufficient to drive detachments (Fig. 4b-c, e-f).

**Fig. 4:**
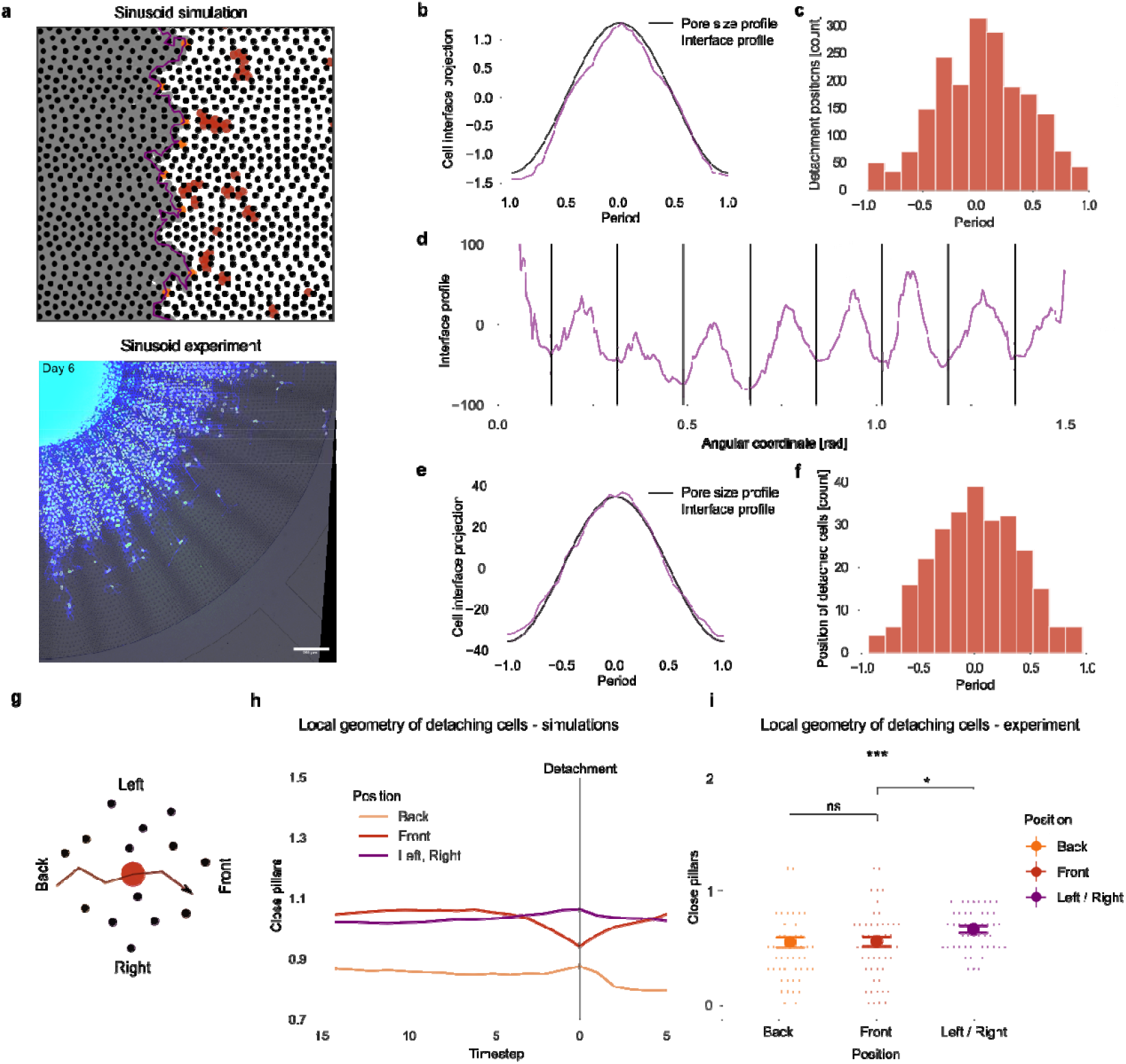
Predicting the statistics of detachments as a function of global and local heterogeneity. **a**. Comparison of simulations (top) and experiments (bottom) using a longitudinal sinusoidal pillar pattern that introduces periodic variation in lateral pore size. Cells invade with forming regularly spaced finger-like protrusions. Simulated cells are shown in grey, pillar in black, protrusion tips in orange, cell interface in magenta, detached cells in red. In experiment, cell bodies express cytoplasmic mCherry (blue) and nuclear H2B-GFP (green). **b**. Simulated interface profile averaged over all timepoints and sinusoidal periods (n=10 simulations), showing alignment of protrusions with regions of lower pillar density. Shaded area indicates standard error. **c**. Spatial distribution of detachment events in simulations across all timepoints and periods of sinusoid profile showing a strong preference for detachment at low-density regions. **d**. Interface profile in experiment in sinusoidal geometry, averaged over all timepoints and plotted in spherical coordinates (n=4 experiments). **e-f**. Average interface profile (e, purple) and corresponding positions of recently detached cells (f, red bars), showing strong agreement with simulation predictions. **g**. Schematic of the method used to quantify local anisotropy in pillar geometry. At each spatial location, surrounding pillars are grouped into 4 quadrants – front, back, left and right. **h**. In simulations, cells approaching detachment exhibit a decrease in front-facing pillar density and an increase in side-facing density, leading to detachments in locally anisotropic regions (time=0 at detachment, n>10000 detachment events from 40 simulations, plot shows mean ± SEM). **i**. Experimental measurements of local pillar density at the time of detachment (n=68 detaching cells from 7 experiments). Cells detach more often in regions with increased side confinement, as simulations predicted. Dot plot shows mean ± SEM. Statistical comparisons (Wilcoxon test): Back vs. Left/Right, p = 0.0045; Front vs. Left/Right, p = 0.018; Back vs. Front, p = 0.74. Together, these results show that both global pattern-induced interface shaping and local anisotropic confinement jointly predict where cells detach from the invading front.

Finally, we asked whether additional local geometrical features predict detachment beyond global integrated heterogeneity. Indeed, as discussed above, repeated simulations on the same disordered geometry revealed “hot-spots” where more detachments occurred. Given that sinusoidal patterns aligned with the invasion axis did not yield detachments (Supplementary Fig. 1d-e, 3c-d), whereas perpendicular sinusoidal did (Fig. 4a-c), we quantified metrics of local pillar anisotropy relative to the invasion direction (front, back, sides; Fig. 4g, Supplementary Fig. 4e-f). In simulations, regions with locally anisotropic free space (i.e. fewer pillars parallel to invasion compared to perpendicular) displayed a small but significant correlation with detachments (R^2^=0.1, p < 10^-4^). When combined with the long-term trend of roughness-induced increases in detachment, predictive power increased substantially (R^2^=0.64, p < 10^-4^, Supplementary Fig. 4e). Consistently, individual simulated cell trajectories showed increased pillar anisotropy specifically at detachment events (Fig. 4h). To test this experimentally, we quantified local pillar anisotropy across n=68 detaching cell trajectories. We found indeed significantly more pillars oriented perpendicular than parallel to the invasion direction at the time of detachment (Fig. 4i). Together, these results demonstrate that disordered environments promote detachments primarily via globally increased interfacial roughening and finger-like protrusions, but that local substrate anisotropy further modulates the likelihood of cell detachment.

## Discussion

Our results demonstrate that environmental heterogeneity is a key driver of cancer cell dissemination that the geometry of the collective invasion front critically determines detachment events. We find that substrate heterogeneity drives detachments from the bulk through two complementary mechanisms. First, as cells invade disordered environments, the interface gradually roughens, effectively integrating heterogeneity over time. This appears to be a generic effect, since the roughening dynamics map onto the KPZq universality class of growing interfaces. After a critical level of roughness is reached, cells at the tips of protrusions have few enough neighbors to detach from the collective. Second, this global permissive effect is compounded by local substrate geometry, where anisotropic pillar arrangements selectively increase detachment probability at specific sites. Thus, disorder influences invasion both globally and locally, providing a robust physical framework that may generalize to other collective cell systems beyond cancer.

Although our study was conducted in a controlled *in vitro* setting, where environmental geometry could be finely tuned, the next challenge is to relate our findings to more realistic *in vivo* situations^62^. For instance, different organs exhibit different extracellular matrix architectures^63^, which often fall between fully ordered and fully disordered states, although this spectrum has not been systematically quantified. Generalizing our simulations to 3D fibrous environments could provide new insight into how tissue organization shapes invasion through environment-tumor cross-talk. In line with that idea, quantitative imaging of human tumor samples revealed that spatially disordered or tilted stromal architectures can be negative prognostic factors for clinical outcome^64,65^. Another natural extension of the model would be to incorporate remodeling of the environment due to mechanical forces exerted by cell migration^66^, which could create feedbacks between cells and their microenvironment that further influence invasion dynamics. We anticipate that combining microengineered models, minimal simulations, and quantitative pathology could yield a predictive framework for how stromal heterogeneity impacts tumor progression.

Using an experimentally controlled bottom-up approach, we introduce stromal heterogeneity as a potential driver of malignant progression. In analogy to organismal evolution, where heterogenous habitats drive genotypic diversity, a heterogenous tumor stroma might shift tumor cells into novel phenotypic states (in our case, the single cell rather than the collective state). Although these states are transient phenotypes without heritable traits, they may later become consolidated^28^ by epigenetic reprogramming and ultimately genetic accommodation. Our findings therefore highlight a potential route by which non-genetic environmental disorder can generate phenotypic diversity that later becomes fixed during tumor evolution.

## Acknowledgments

The authors thank all the members of the Hannezo and Sixt groups for fruitful discussions and Takuya Kato for advices on the experimental set-up. We are also thankful for support from the Scientific Service Units (SSU) of IST Austria through resources provided by the Imaging and Optics Facility (IOF) and the Lab Support Facility (LSF). Z.D. has received funding from the Doctoral Programme of the Austrian Academy of Sciences (OeAW): grant agreement 26360. This work was supported by the European Research Council (grant ERC-SyG 101071793 to M.S).

## Author contributions

S.T., Z.D., M.S., and E.H. designed the research. S.T. performed the experiments: S.T., Z. D. and Ju.M. performed data and image analysis; Z.D. and Ju. M. performed simulations; Ja.M. supported microfluidic engineering; E.S. provided cellular model systems. M.S. and E.H. supervised the study; S. T., Z.D., M.S. and E.H. wrote the manuscript, with input from all authors.

## Methods

### Experiments

#### Cell Culture

A431 cells expressing mCherry were grown in DMEM (Gibco, 10569010) containing phenol red, high glucose, GlutaMAX, pyruvate, 10% fetal calf serum (FBS, Gibco, 10270106), non-essential aminoacids (Gibco, 11140050), penicillin (50U/ml) and streptomycin (50μg/ml, Gibco 15070063) in humidified incubators at 37°C with 5% CO_2_. EPCad^-/-^ A431 cells were made as described in Labernadie et al. 2017^67^. Cells were regularly tested for mycoplasma contamination and preserved in growth medium containing 5% (v/v) DMSO in cryogenic conditions for longer periods. Cells were transfected with lentivirus containing H2B-GFP construct and selected for blasticidin resistance and then later sorted for GFP signal, regularly.

#### Device Fabrication

Patterns were generated with Python script in SVG format. Designs of photomasks were completed in (Inkscape, v0.92) in layers for each confinement. SVG files were converted to AutoCAD DXF R14 files, further converted to Gerber files (LinkCAD) and finally printed on chrome photomasks at a resolution of 1 μm (JD Photo Data & Photo Tools). For high-resolution structures, wafers were first coated with a sub-micrometre layer of SU-8 as a sticky substrate for later SU-8 structures. We used GM-1040 SU-8 photoresist (Gersteltec) at 5,000 rpm for 2 min with flood UV exposure. The feature layer was then applied with SU8-2005 (Microchem). According to the datasheet, a UV-absorbing glass plate and higher exposure were used to make sharp features. PL-360-LP from Omega Optical and 550 mJ/cm^2^ with an EVG mercury lampbased mask aligner gave good results in the 3–9-μm-thickness range. The thick (25 μm) loading channels were made with Microchem SU8-3025 using the EVG mask aligner according to the Microchem data sheet without the PL-360-LP plate. The master wafers underwent vapour silanization for 1 h with (1H, 1H, 2H, 2H-perfluorooctyl) trichlorosilane. The final results were single hydrophobic master wafers containing several aligned layers. Polydimethylsiloxane (PDMS) devices were made with Sylgard 184 Elastomer Kit (Dow Corning) using a ratio of 10:1. The PDMS was mixed in a Thinky mixing machine and then poured on the wafer in a 3D printed tray and further degassed several times in a vacuum chamber. The devices were baked at 80 °C overnight. The hardened devices were carefully peeled off the mould and cut into single de-vices using a razor blade. Holes were punched with a custom arbor press or with skin biopsy puncture tools (1 mm) for the central hole, and sticky tape was used to remove debris. Glass coverslips were sonicated for 5 min in 100% isopropanol and blown dry. Devices were bonded to the glass slides by oxygen plasma then sealed together at 85 °C for 1 h to make the bond permanent.

#### Invasion Assays

Microfabricated devices were inserted into 12-well cell culture plates from the bottom side and glued with molten paraffin. Devices were filled with fibronectin (F1141, Sigma) solution (diluted in 1:5 from 1mg/ml stock solution in PBS) using a syringe pump, sealed and incubated for 3h in cell culture growth conditions. Devices were later flushed with 100 μl cell culture growth medium with syringe pumps. Debris were removed from the center hole, either spheroids of cancer cells (generated with hanging drops) were dropped or a pipette tip filled with medium was inserted into the hole, where 1.5 μl of the cell suspension was filled. Cells were incubated until they sank into the hole and imaged at later days. Wells were checked for contamination under microscope.

#### 2D Spread Assay

A431 cells were mixed with cytoplasmic mCherry reporter expressing A431 cells (10:1) and assembled into 1000 cell spheroids in hanging drops overnight. Glass cover slips were coated with fibronectin (F1141, Sigma, diluted in 1:40 from 1mg/ml stock solution in PBS) for 1h and then washed with PBS. Subsequently, chambers were filled with cell culture media and spheroids were placed to adhere on the substrate. Cells were observed with fluorescence and DIC microscopy.

#### Scratch Assay

In order to observe invasion on non-confined 2D surface with an initial straight interface, cells in suspension were placed and grown to full confluence on fibronectin coated glass substrates (see 2D spread assay above). Then, approximately at the center of the cover slip, cells were scratched with a rubber cell scraped and detached cells were aspirated and washed away with cell culture media. Cells were observed with fluorescence and DIC microscopy.

#### Imaging

Cells were either imaged once a day at room conditions or in cell culture conditions for timelapse imaging (10 min interval) for several days (typically 3-4 days). Invasion was recorded with an inverted wide-field Nikon Eclipse Ti-2 microscope equipped with a Lumencor light source (475 nm for GFP and 594 nm for mCherry) and an incubation chamber, heated stage and CO_2_ mixer (Ibidi). Generally, the fluorescence imaging was accompanied with light microscopy (DIC) imaging. For long term imaging, phenol red color was checked and the temperature was monitored with a temperature sensor. Custom made CNC milled box and a pair 3D printed plugs were used to prevent CO_2_ and humidity leak.

#### Collagen Assay

A collagen mixture was created with a final collagen concentration of 1.7 mg/ml by mixing bovine collagen (PureCol, Advanced BioMatrix) in 1x minimum essential Eagle (Sigma) and 0.4% sodium bicarbonate solution (Sigma Aldrich). Mixture was checked for a pale pink color to ensure for correct pH for polymerization. 20 μl of the collagen mixture was mixed with 9 μl of the growth medium and mixed gently. Subsequently, 1 μl of the spheroid containing growth medium was added to the mixture and mixed by stirring with the tip. Final mixture was gently placed on a glass coverslip, which was previously glued under a 60 mm dish with molten paraffin. The dish was flipped to let the collagen drop hang from the glass coverslip, whereas the plastic lid was filled with droplets of PBS to keep the dish humidified inside the incubator. Collagen drop was let to polymerize in the incubator for 45-60 min under cell culture conditions. Before imaging, PBS was removed and the dish was flipped again and medium was added to the chamber with the collagen drops. Invasion into the collagen was imaged with light microscopy (DIC) over several days under cell culture conditions.

#### Lentivirus Production

Lentivirus production was performed as described previously^68^. In brief, LX-293 HEK cells (Clontech) were co-transfected with LentiCRISPRv1 or LentiCRISPRv2, packaging-ps-PAX2 (Addgene no. 12260) and pCMV-VSV-G envelope plasmids using Lipofectamine 2000 (ThermoFisher Scientific) as recommended by the manufacturer, and cells were resuspended in the growth medium 1 day before transfection. Supernatant was collected after 72 h and stored at −80 °C.

### Image processing and experimental analysis

#### Image Analysis

FiJi was used for image processing, including using the TrackMate plugin for manual cell tracking^69,70^. Proprietary ND2 files from Nikon NIS Elements were converted using Bio-Formats plugin^71^ and the physically calibrated (pixel size and imaging interval). Images were aligned and processed into TIFF and HDF files. Most of the analysis processes were automated with ImageJ macros. Cell nuclei (H2B-GFP, green) and the remaining cell body (mCherry, blue) and nuclear H2B-GFP (green) were initially segmented using Ilastik and then particle analysis feature was used to quantify invasion^72^. Cells were tracked semi-automatically with TrackMate, tracks were corrected manually. Dividing and dying cells were excluded, all gaps were filled with interpolation. Final images were processed with automated scripts for display and exported into PNG. In images, cells were considered detached, when their cell bodies were not considered touching the bulk. Finger measurements were performed manually by measuring the distance from the tip to the finger baseline. Detachments were selected over a wide range of recorded experiments. The pore size of the detaching cell during detachment vs. the pore size in front of the cell, which is being detached from, was measured manually with an interactive macro.

#### Experimental data analysis

The cell interface in experiment was obtained by identifying the nuclei positions of cells at the interface, for better comparison with simulations, which do not model the cell bodies (Supplementary Fig. 4a). Identification of cells at interface was done by first running Delaunay triangulation of all cell nuclei in each timestep and subsequently removing the vertices, which cross through the segmented interface of bulk cell bodies (blue line in Supplementary Fig. 4a). This way we could efficiently find all cells, which are in contact with the very rough and irregular bulk interface. Before using the interface positions for analyzing the roughness exponents, we had to remove the overall tilts in the interface present in some experiments (see left panel in Fig. 1d, Supplementary Fig. 5a). This was done by removing long-wavelength Fourier modes (> 500 μm). The analysis of the local pillar geometry around the detaching cells (Fig. 4i) was performed by identifying all close pillar in a given radius around a the x,y position of the detaching cells and dividing them into 4 quadrants (front, back, left right, see Fig. 4g, the center of the front quadrant being the orientation of the detaching cell). Multiple radii were tested, all resulting in qualitatively similar result (final value: r_max_= 27 μm, i.e three times the pore size). The identified pillars in each quadrant were subsequently weighted by their distance from the reference position (r_max_-r)/r_max_ and summed.

#### Statistics and Data Visualization

All statistics for experimental data were performed in *R (4*.*0*.*3)* and plotted using *ggplot2* package (R Core Team 2020; Wickham 2016:2). Tracking data was analyzed with *TraXpert*, an inhouse developed particle tracking analysis application. Directionality data was analyzed with *circular* package and circular statistics^73,74^. Minor cosmetic adjustments were performed in *Inkscape*, without affecting the data points or the interpretation of the plots. Figures were assembled in *Inkscape* and exported into high resolution lossless PNG images.

### Numerical model

We implemented two dimensional Brownian dynamics simulations as a minimal model of collective cell migration^44–47^ through complex environments (Fig. 2a). In this framework, each cell is represented as a soft self-propelled particle (approximating its nucleus^41^) moving on a substrate. The overdamped equation of motion for each particle is given by:

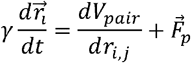

where γ is the friction coefficient, 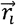 the particle coordinates, _pair_ the pairwise interaction poten-tial, and *r*_*i*,*j*_ the distance between particles. The active self-propulsion force 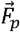 is given by

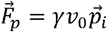

Where *v*_0_ is the intrinsic propulsion speed and 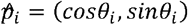 defines the orientation of parti-cle. In the absence of interactions, this force yields an average cell speed of *v*_0_.

The polarity of migrating cells is subject to rotational diffusion and an external biasing torque *τ*_bias_, which directs cell migration away from the bulk (see below for discussion of its origin). The orientation dynamics are given by

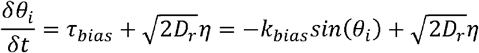

where *D*^*r*^ is the rotational diffusion coefficient, a unit-variance Gaussian random variable, and *k*_bias_ controls the strength of the bias torque. We define *θ*_*i*_ = 0as the invasion direction (to the right in all simulations shown). To validate the numerical implementation, we computed the mean square displacement of a single active Brownian particle without bias torque^75^. The observed crossover from ballistic to diffusive motion was consistent with the used value of *D*_*r*_.

To describe cell-cell interactions, we assume a simple pairwise potential consisting of short-range repulsion and intermediate-range attraction, as previously described^44,45,76^. The potential takes the form

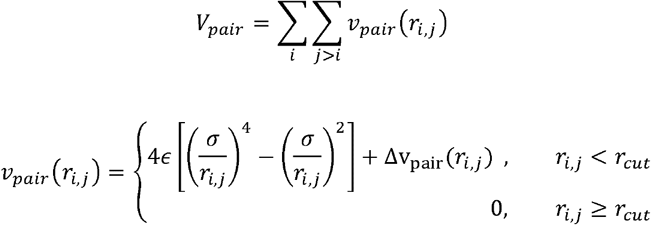

where *r*_*i*,j_ is the distance between particles *i* and *j*, □ is the depth of the potential well, σ is the characteristic interaction distance, and *r*_*cut*_ is the cutoff distance. The potential is shifted by a linear correction term Δ*v*_*pair*_, ensuring that both the potential and its derivative smoothly vanish at the cutoff 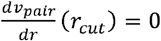.

To model the structured environments (pillar geometries), we introduced stationary particles into the simulation box, which contribute to the total pair potential. The cell-pillar interaction was truncated at its minimum 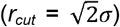 so that only repulsive interactions remained, consistent with the absence of strong experimental evidence for stable cell-pillar adhesion. Importantly, a central assumption of our minimal modeling approach is that intrinsic cellular parameters (e.g., motility, adhesion strength) are held constant across conditions; thus the observed differences in detachment dynamics arise solely from variations in substrate heterogeneity.

### Details of the simulation setup

We used the HOOMD-blue v3 library to perform numerical simulations^77^. The equations of motion were integrated using the Euler-Maruyama scheme with a timestep of *dt* =0.001. All simulations were conducted in two dimensions.

Simulation parameters and results are reported in dimensionless units. For convenient comparison with experiments, lengths are expressed in units of cell-cell equilibrium distance *L* (defined as the interaction minimum 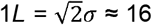 um) and time in units of = 10^3^ s. Simulation output was saved every 600 frames, corresponding to the smallest experimental timestep (10 minutes).

Unless stated otherwise, the simulation parameters used in the model are summarized in Supplementary Table 1:

**Supplementary Table 1:** Summary of the simulation parameters.

| Parameter | Notation | Chosen value<br>(simulation units) | Parameter fitting |
| --- | --- | --- | --- |
| Friction coefficient | $\gamma$ | 1 | Fixed parameter, non-dimensionalizes timescales |
| Attraction cutoff | $r_{cut}$ | 1.55 $L$ (1 $L$ for cell-pillar interaction) | Fixed parameter (on the order of cell size to avoid next-neighbor interactions) |
| Self-propulsion speed | $v_0$ | 0.43 $L/\tau$ | Fitted to correspond to mean single-cell speed $6.9 \cdot 10^{-3} \mu\text{m/s}$ inside the pillar pattern (Supplementary Fig. 5a,b) |
| Rotational diffusion | $D_r$ | 0.7 $\tau^{-1}$ | Fitted according to MSAD of experimental trajectories of detached cells (Supplementary Fig. 5c) |
| Bias torque | $k_{bias}$ | 0.75 $\text{rad}/\tau$ | Fitted according to the drift of experimental trajectories of detached cells (Supplementary Fig. 5d) |
| Cell fluidity number | $\gamma v_0/\epsilon$ | 0.52 | Constrained by cell confluency while allowing occasional bulk cell rearrangements (Supplementary Fig. 2b, Supplementary Movie 4, Supplementary Table 3) |
| Pore size | $L_p$ | 0.88 - 1.3 $L$ | Free parameter (on the order of cell size; lower bound ensures cells can pass through pores individually) |
| Pore heterogeneity | $\mathcal{H}$ | $\mu = b = 0.1 \cdot L_p - 0.5 \cdot L_p$ | Defined by parameters of Laplace distribution; upper bound taken from disordered pattern in experiments |

Before doing a systematic parameter search, we first constrained parameters to reasonable ranges and conducted non-dimensional analysis to identify the key dimensionless ratios. The fluidity of a tissue in the absence of obstacles is governed by a dimensionless number relating self-propulsion to cell-cell interaction strength (*v*_*0*_γ*/ ϵ* )^46,78,79^. If the ratio is too low, cells behave as a solid, with very few neighbor rearrangements. Conversely, above a certain threshold, the tissue fluidizes, as motility-driven rearrangements become possible. However, when considering not only bulk tissue but also a cell interfaces, very large values of this ratio can lead to spontaneous cell detachments in pillar-free environments, resembling a gas^80^. Importantly, experiments with monolayers invading a homogeneous environment support an intermediate value of this ratio, as we observed virtually no detachments, while cells remained able to move relative to one another (Supplementary Fig. 1b, Supplementary Movie 4, Supplementary Table 3).

Furthermore, we leveraged the presence of single detached cells in pillar experiments to constrain several parameters related to single-cell migratory properties. In particular, by calculating instantaneous (x,y) displacements and the mean squared angular displacement (MSAD)^81^ of detached cells in both simulations and experiments, we could strongly constrain the values of self-propulsion speed (v_0_) and particle persistence length (v_0_/D_r_) (Supplementary Fig. 5b-d). The value of external bias (*k*_*bias*_) was fixed by analyzing the radial drift in trajectories of detached cells in experiment and comparing it to simulations (Supplementary Fig. 5e). Notably, we detected strong biased migration even in single cells far from the leading edge, indicating that this bias does not simply arise from pushing forces due to bulk proliferation. Given previous reports of self-generated gradients guiding long-range migration^82,83^, such mechanisms are compatible with this bias, although we chose to remain agnostic about its origin in the simulations. Finally, we performed an additional sanity check by comparing the mean squared displacement (MSD) of single detached cells in simulations and experiments, which allowed us to validate the chosen combination of parameters *D*_*r*_, *v*_*0*_, and *k*_*bias*_. The final parameter set reproduced the experimental MSD curve most closely, yielding the highest coefficient of determination (pseudo-R^2^ = 0.89), particularly at long times (Supplementary Fig. 5f).

The simulations of migrating cells used for comparative statistics across various pillar geometries and parameters (Fig. 2b, Supplementary Fig. 2ab, Supplementary Fig. 5, Supplementary Table 2) were initialized in a rectangular box of size *Lx*=*Ly*=425 *L*. The left portion of the box was filled with cells in hexagonal packing, while the right side contained the pillar geometry. Simulations used for close temporal comparison with experiments (Fig. 2c-e, Fig. 3-4, Supplementary Fig. 3-4) were initiated in a smaller box (Lx=425 *L*, Ly=106 *L*) with new cells added on the left side during the simulation to maintain a stable bulk pressure. Each parameter combina-tion was run with at least 10 repeats. The ordered pillar geometry was constructed as a square lattice, with pillar spacing being twice the effective pillar diameter. Reported results for ordered pattern simulations (e. g. Fig. 2b,c,e and Fig. 3d) are averaged over four orientations of the square lattice (0°, 15°, 30° and 45°, Supplementary Fig. 3b). The disordered pattern was generated from the ordered patterns of all four tilts by shifting each pillar by a random angle and distance following a Laplace distribution from its original position. The fully disordered pattern corresponded to Laplace distribution parameters μ = *b =* 0.5 * L_p_.

#### Simulation analysis

The initial time required for cells to accumulate at the front of the pillar pattern and enter it was excluded from the analysis. Unless stated otherwise, simulation frames were analyzed at a frequency of 6 τ after the start of the invasion. Simulations were run until bulk cells reached the end of the pillar pattern. All simulation analyses were performed in Python.

The overall number of detached cells (Fig. 2b,d, Fig. 3a, Supplementary Fig. 2a,b,d, Supplementary Fig. 3d) was determined by distance-based clustering of cells, using *r*_*cut*_ as the natural clustering cutoff from a physical perspective, and summing the number of cells within the pattern that were not connected to the bulk at a given timepoint. The identification of pattern locations with high detachment probability and instantaneous detachments (Fig. 2c, Fig. 3c-d,f-g, Fig. 4c,h, Supplementary Fig. 2e, Supplementary Fig. 3g, Supplementary Fig. 4c-e) was performed in two steps. First, the same clustering analysis was applied to each simulation frame. Second, a cell was considered detached if it remained separated from the bulk (largest cluster) for at least 5 consecutive frames. For each detachment event, we recorded the x,y coordinates, cell ID, and timestep for further analyses, such as relating detaching events to interface properties (Fig. 3c,d) or local pillar positions (Fig. 4d).

The shape of the cellular interface (eg. Fig. 2d,e, Fig. 3a, Fig. 4a) was analyzed by first computing the Delaunay triangulation (convex hull) of cellular positions in each simulation frame. The Delaunay triangulation was then filtered using the alpha shape algorithm^84^ (Python alphashape library) to determine the concave hull of the cellular interface (alpha = 0.6). The positions of interface cells were subsequently used in the roughness scaling analysis (Fig. 2e, Supplementary Fig. 3c,e,g) and in the analysis of interface shape, perimeter and curvature (Fig. 3a,c,d,h, Fig. 4b, Supplementary Fig. 3a). The local maxima and minima of the cell interface were identified using the argrelextrema function in SciPy library^85^. Transient protrusions on the order of cell size (protrusion length = average distance to next local minimum < 1 *L*) were filtered out.

The interface roughness (Fig. 2e, Supplementary Fig. S3e,g) was quantified using the height function *h(y)*,which represents the level of invasion of cells at different points along the interface (polar coordinates were used for pillar pattern experiments). We analyzed the temporal increase in roughness by computing the interface width *w(l*,*t)*, which measures the root-mean-square deviation of the interface from its mean value, defined as:

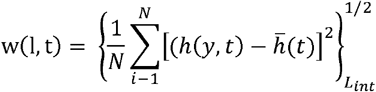

where *L*_*int*_=1000 μm is the length of the interface contour. These fluctuations grow over time following a power law scaling,w(l,t) ∼*t β*, with a characteristic critical growth exponent *β* ^31^.

To quantify the spatial roughness of the invasion front (Supplementary Fig. S3b), we computed the height–height correlation function of the interface profile in simulations. The function is defined as:

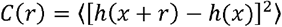

Where *h(x)* denotes the height of the interface at lateral position *x*, and ⟨⋅⟩ represents an average over all positions *x* and multiple simulation runs. For self-affine interfaces, this function exhibits power-law scaling with distance *r*:

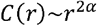

We computed *c*(*r)* from late-time snapshots of the invasion front, after roughness had reached a statistically stationary regime. The roughness exponent *α* was extracted by fitting the log–log plot of *c*(*r*) versus *r* over the intermediate scaling range, avoiding short-range discretization artihomogeneous and disordered environments. The resulting values of *α* provide an independent facts and long-range saturation effects. We repeated this analysis for simulations conducted in estimate of the interface roughness exponent, complementary to the growth exponent *β* obtained from time-dependent roughness evolution. To test for Family–Vicsek scaling (Supplementary Fig. S3c), we performed simulations for multiple lateral system sizes *L*. For each *L*, we ran and averaged ∼20 independent simulations; in homogeneous environments these differed only by stochastic cell dynamics, whereas in disordered environments they additionally differed by independently generated random pillar patterns. We then rescaled roughness and time as *W*/*L α* and *W*/*L β*, with independently measured exponents (*α* = 0.55, *β* = 0.33 in homogeneous; *α*= 0.70, *β* = 0.55 in disordered environments) and the dynamic exponent defined as *z* = *α*/ *β*. Collapse of the rescaled curves across system sizes was taken as evidence that the observed dynamics fall into the corresponding universality class.

The analysis of local pillar heterogeneity (Fig. 4h, Supplementary Fig. 4e,f) was performed by identifying all neighboring pillars within a radius r_max_= 3.18 *L* (three times average pore size) around a given (x,y) position in the disordered pattern. These pillars were then classified into four quadrants (front, back, left right; Fig. 4g). The pillars in each quadrant were weighted by their distance from the reference position using the factor (r_max_ - r)/r_max_ and the weights were then summed.

## Supplementary Figures

**Supplementary Fig. 1:**
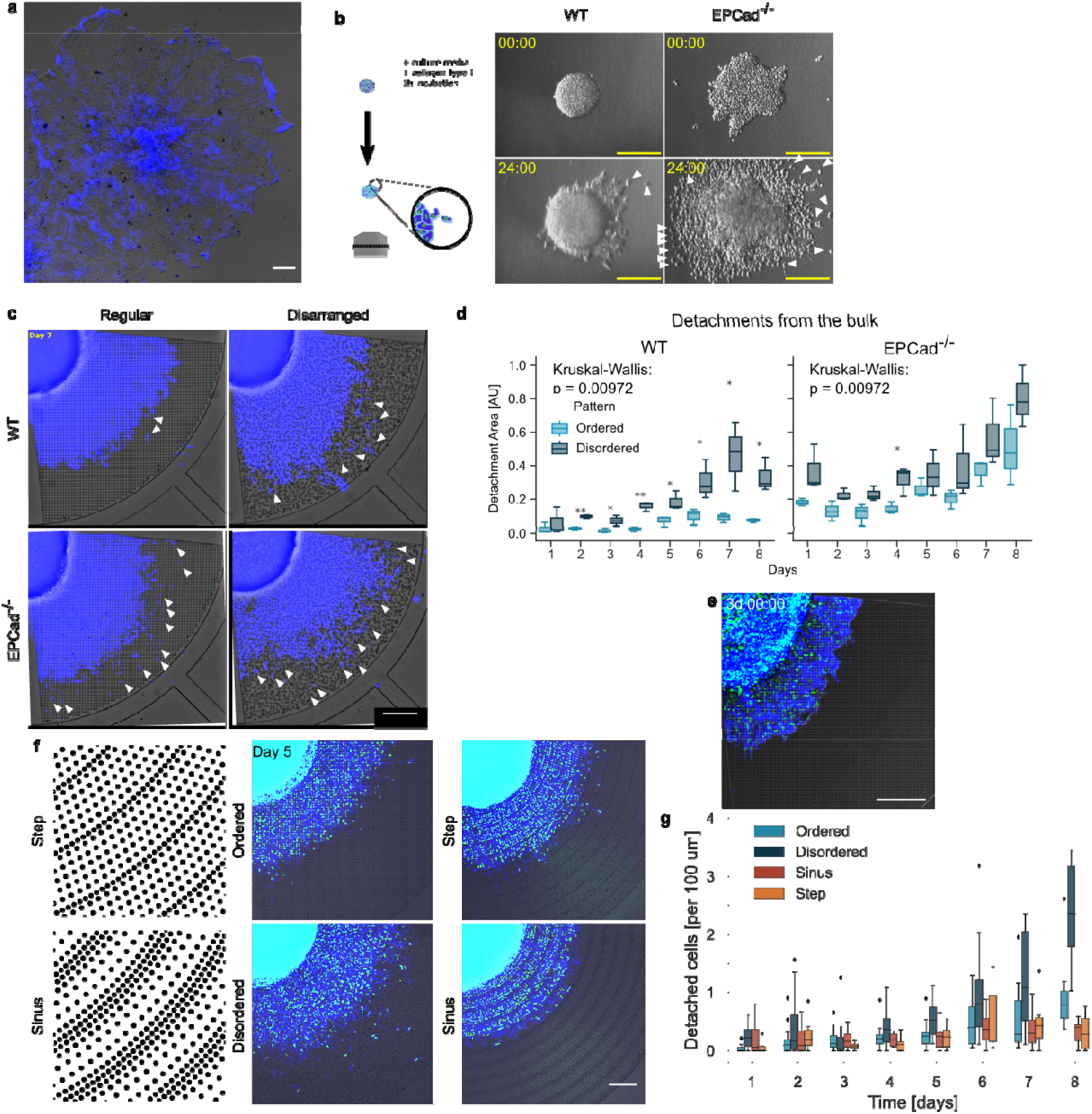
Invasion dynamics of cells in different environments. **a**. 2D invastion assay of WT A431 cells with and without mCherry reporter with 1:10 mix respectively. **b**. Collagen invasion assay of WT and E-,P-Cad^-/-^ spheroids imaged with brightfield microscopy. **c**. Detachments of the WT and E-,P-Cad^-/-^ cells invading into ordered and disordered pillar patterns. White arrows indicate individual detached cells or cell clusters. **d**. Detached cell area in the respective pillar patterns for A431 WT and E-,P-CadKO cells. Scale bars: 100 μm, box and whiskers plot displays median, 1^st^ and 3^rd^ quartiles (hinges) and bars display 1.5 interquartile range from the hinges, n=3 experiments per genotype and pattern followed over 8 days. **e-f**. Cell invasion assays into smaller (6 μm) sized ordered pillar pattern (d) and stepwise or sinusoidal pattern (e). **g**. Quantified number of detachments for each pillar patterns, where box and whiskers plot displays median, 1^st^ and 3^rd^ quartiles (hinges) and bars display 1.5 interquartile range from the hinges, n=3 experiments per genotype and pattern followed over 8 days. Statistical comparisons (Mann–Whitney U): Ordered vs. Sinusoid, p = 0.61; Ordered vs. Stepwise, p = 0.88). All scale bars: 200μm.

**Supplementary Figure 2:**
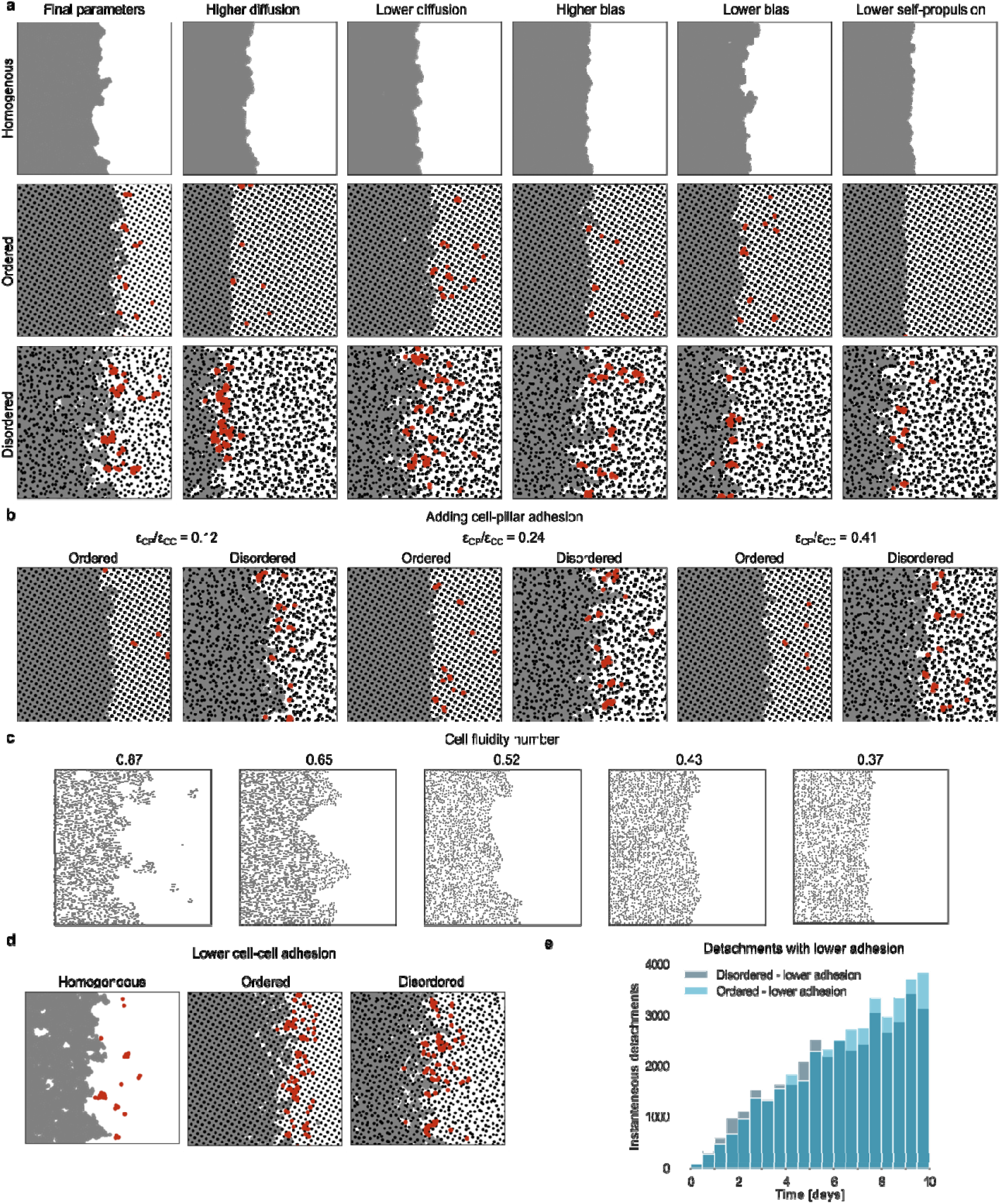
Phase diagram of invasion dynamics for different model parameters. **a**. Temporal increase in detachment rate is a robust phenomenon present both in ordered and disordered pillar patterns, as well as for different simulation parameters. The simulation snapshots show a representative state of the systems at day 6 (cells are shown as grey and pillar as black beads, detached cells are highlighted in red), with the best-fit set of simulation parameters used in the main text (left column, see also Supplementary Table 1). Each subsequent column shows the effect of changing a single simulation parameter (*D*_*r*_ = 1 ^*-1*^, D_r_ = 0.5 ^*-1*^, *k*_*bias*_ = 0.5 *rad/*, k_bias_ = 1 *rad/*, *v*_*0*_ *=* 0.24 *L/* respectively). **b**. Snapshots of simulation od cells in ordered and disordered environments at day 6 (cells are shown as grey and pillar as black beads, detached cells are highlighted in red), for different values of cell-pillar adhesion (r_cut_=1.55 *L)*. **c**. Snapshots of simulation of cells in homogeneous environments (grey beads shown with smaller diameter for clarity, day 6), for different values of the fluidity number. High fluidity number (left) leads to numerous detachments, something not observed in wild-type experiments, while low fluidity (right) leads to a solid-like rheology without any re-arrangements on the time scale of the simulations (also not observed). **d**. Snapshots from simulations of cells with lower cell-cell adhesion, corresponding to E-,P-Cad^-/-^ cells (fluidity 0.87, cells are shown as grey and pillar as black beads, detached cells are highlighted in red). This leads to detachments even in the absence of any pillars, i.e. to a much weaker sensitivity to the environment, as seen experimentally. **e**. Quantification for the increase of detachments when lowering the cell-cell adhesion, for either ordered or disordered pillar patterns, showing that the difference between ordered and disordered pattern disappears (T-test: p=0.47) compared to intermediate adhesion/fluidity.

**Supplementary Figure 3:**
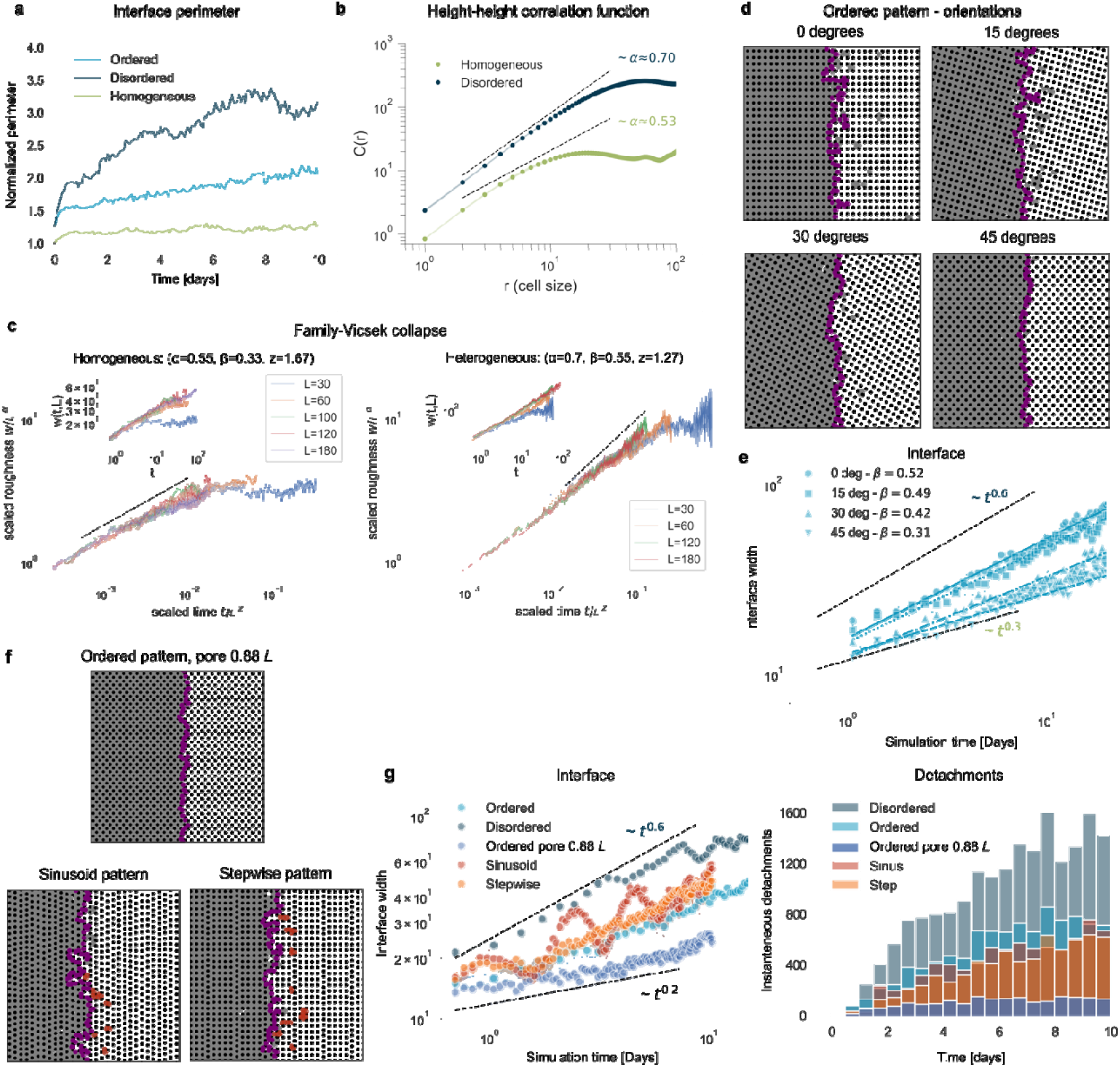
Universality class identification and sensitivity analysis of simulation outputs for different pillar patterns. **a**. Temporal evolution of the interface perimeter, quantified in simulations (n=40 simulations, the shaded intervals indicate standard error). **b**. Height-height correlation function of the invasion front in simulations in steady-state snapshots of the interface (day 25) to estimate the roughness exponent. Shaded intervals represent standard error across the analyzed snapshots. Fits were performed over the intermediate scaling range (1 < r < 12). **c** Family–Vicsek scaling collapse of interface roughness for homogeneous (left) and disordered (right) environments. When the original curves for different *L* (insets) are rescaled according to the Family–Vicsek relation / vs. /, with independently measured exponents, they collapse onto a single universal function, confirming assignment to the KPZ and KPZq universality classes, respectively. **d**. Differences in invasion behavior with different orientations of regular pattern (snapshots of day 4, cells are shown as grey and pillar as black beads, magenta line represents cell interface). We used the average results of the 4 shown orientations for comparing regular pattern with other geometries, given that invasion is radial in experiments. **e**. Interfacial roughness and roughness exponents in 4 orientations of regular patterns (corresponding to b) (n=10 simulations each, the shaded intervals correspond to standard error). **f**. Simulation snapshots of invasion in sinusoid and stepwise pillar patterns, exhibiting lower or comparable detachments, as in ordered pattern, as seen experimentally in Supplementary Fig 1e-g. Cells are represented as grey and pillars as black beads, detached cells are marked red, cell interface in magenta, day 4. **g**. Quantifications of interface width and detachment numbers in simulations of panel d, showing that it follows a stepwise trend corresponding to the pillar pattern, and that some cells tend to reattach to the bulk resulting in lower overall number of detachments.

**Supplementary Figure 4:**
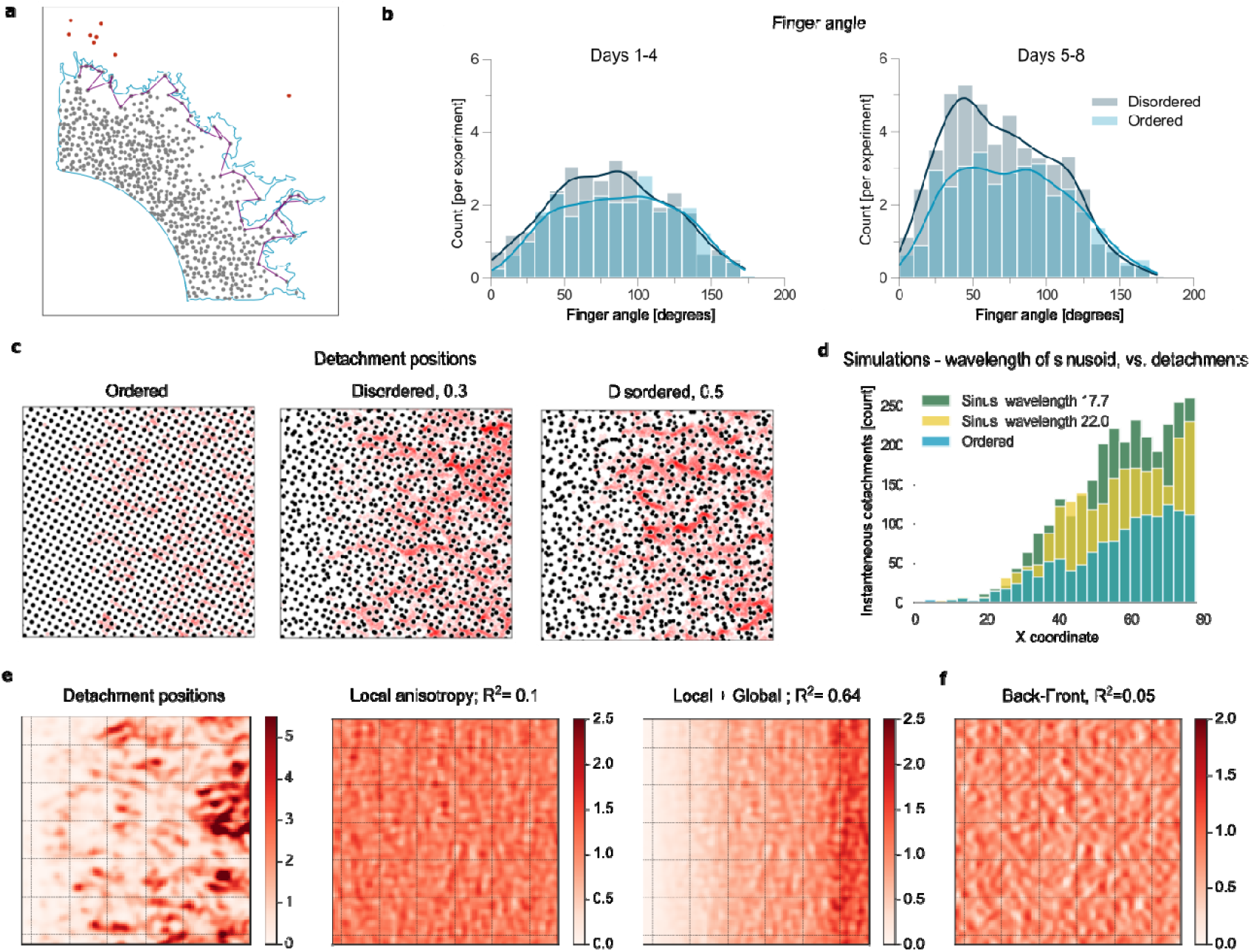
Detachments statistics in data and simulations. **a**. Analysis of interface roughness in experiments. The blue line represents the segmented outline of cell bodies and magenta line connects the cell nuclei along the interface, which are used for subsequent roughness and interface curvature analysis (see Methods). **b**. Difference between the finger angle (defined as the angle between the tip of finger and neighboring local minima, with small angles thus corresponding to sharp fingers, see Fig. 3e) of the protrusions in ordered and disordered patterns, which becomes more pronounced in later timepoints (n=16 experiments for regular and n=18 for disordered pattern). **c**. Distribution of detachment events in simulations with increasing disorder in pillar positions (color coded in red for increasing probability). **d**. Temporal increase of detachments in perpendicular sinusoid simulations with smaller wavelength (see Fig. 4a). **e**. Heat map of detachment positions compared to the local anisotropy of pillar positions in disordered pattern (Metric: Left + Right – Front, R^2^ = 0.1) and local anisotropy of pillar positions multiplied by the mask, which takes into account the increase of detachments in the direction of invasion (R^2^=0.63). detachments analyzed from n=10 simulations. **f**. Example of a different metric calculated using the same analysis (Back-Front), which exhibited lower spatial correlation with detachment positions.

**Supplementary Figure 5:**
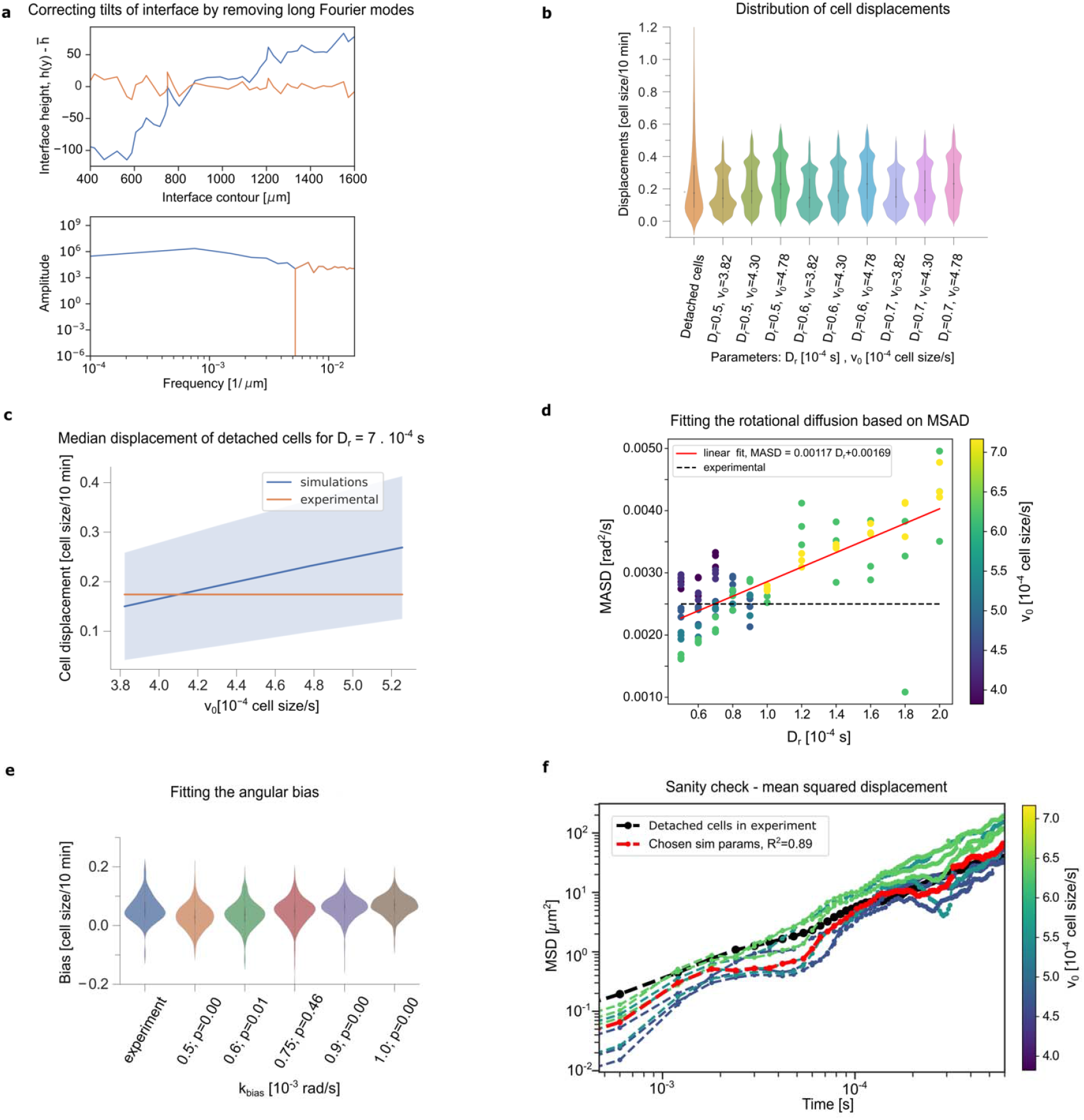
Fitting procedure for different model parameters. **a)** Correction of tilts in the detected experimental interface (blue line) by removing long Fourier modes (orange line). **b-c)** Fitting the active force v_0_ based on displacements of single detached cells in a pillar patterns, respectively (experimental data shown in the first two columns of panel b, orange line in panel c represents the median of distribution of detached cells, n=91 trajectories from n=6 experiments). The displacements do not depend on *D*_*r*_ and grow linearly with v_0_ (further columns in panel b show n=3 simulations of n=100 sparsely added single cells in ordered pillar patterns with pore size 1.06 *L* and different combinations of parameters). By comparing the data to different values of *v*_*0*,_ we constrained *v*_*0*_=0.43 *L/* as the best-fit parameter used in all simulations unless explicitly mentioned **d)** Fitting of the rotational diffusion constant based on MSAD of single detached cells in a regular pillar pattern (color coded according to value of *v*_*0*_, black dashed line represents MSAD of detached cells in experiment). As above, we compared the experimental results from trajectories of detached cells to simulations with different rotational diffusion coefficients D_r_. We found that the best-fit occurred for 0.7 ^*-1*^ (linear fit to simulation data, R^2^=0.93), which we use for all simulations unless explicitly mentioned. **e)** Fitting of the *k*_*bias*_ based on the radial bias in trajectories of single detached cells in a regular pillar pattern (same procedure as above). We found that *k*_*bias*_=0.75 *rad/* reproduced best the experimental data (Kolmogorov-Smirnoff test, p=0.46), which we use for all simulations unless explicitly mentioned. **f)** MSD comparing the detached cells in pillar patterns in experiment and different combinations of simulation parameters *D*_*r*_, *v*_*0*_, and *k*_*bias*_. The final set of parameters chosen according to **b-e** exhibited highest pseudo-R^2^ related to the experimental MSD curve.

**Supplementary Table 2.** Interface roughness of cells invading in ordered and disordered patterns in different parameter combinations (green – exponents from 0.25-0.44, blue – larger than 0.55).

| Pore size | $D_r [\tau^{-1}]$ | Ordered pattern | | | | Disordered pattern | | | |
| --- | --- | --- | --- | --- | --- | --- | --- | --- | --- |
|  |  | fluidity |  |  |  | fluidity |  |  |  |
|  |  | 0.44 | 0.52 | 0.60 | 0.76 | 0.44 | 0.52 | 0.60 | 0.76 |
| 0.88 L | 0.5 | 0.27 | 0.33 | 0.41 | 0.50 | 0.55 | 0.70 | 0.70 | 0.57 |
|  | 1 | 0.19 | 0.36 | 0.43 | 0.44 | 0.45 | 0.65 | 0.72 | 0.71 |
|  | 2 | 0.37 | 0.25 | 0.23 | 0.37 | 0.44 | 0.36 | 0.39 | 0.49 |
| 1.06 L | 0.5 | 0.48 | 0.57 | 0.62 | 0.59 | 0.54 | 0.62 | 0.78 | 0.57 |
|  | 1 | 0.30 | 0.45 | 0.52 | 0.55 | 0.44 | 0.69 | 0.64 | 0.59 |
|  | 2 | 0.34 | 0.23 | 0.28 | 0.42 | 0.35 | 0.32 | 0.40 | 0.43 |
| 1.24 L | 0.5 | 0.32 | 0.32 | 0.38 | 0.38 | 0.57 | 0.66 | 0.80 | 0.49 |
|  | 1 | 0.37 | 0.46 | 0.49 | 0.50 | 0.58 | 0.69 | 0.64 | 0.59 |
|  | 2 | 0.33 | 0.27 | 0.31 | 0.41 | 0.56 | 0.33 | 0.40 | 0.43 |

**Supplementary Table 3:**
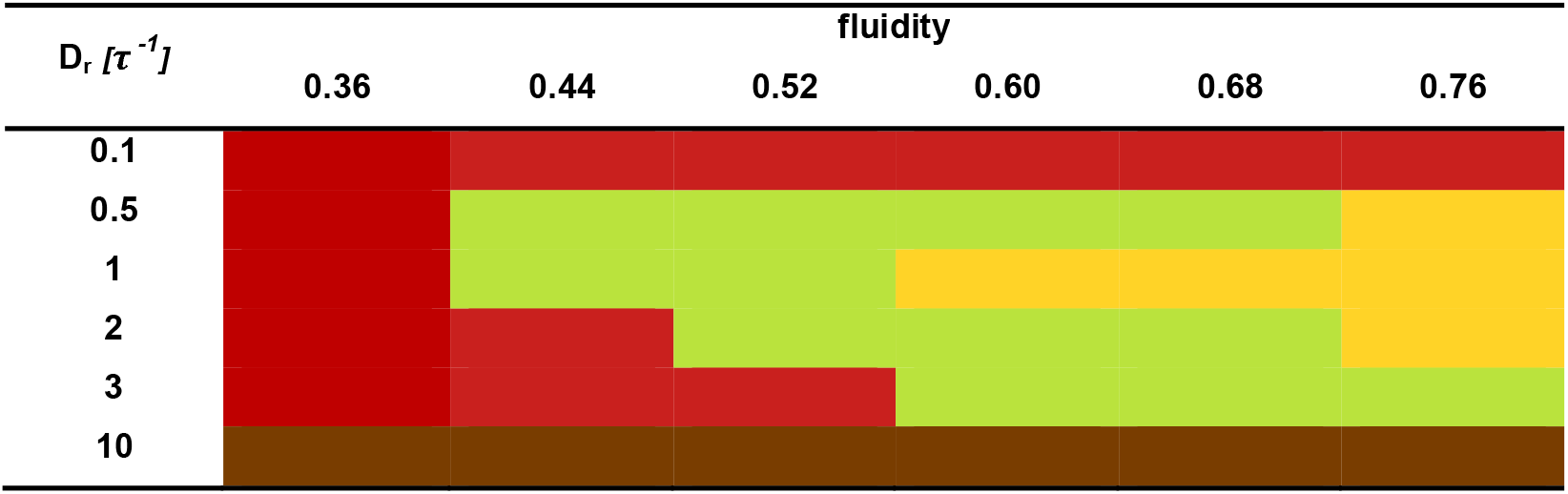
Behavior of the cell bulk in homogeneous environment in simulations with different parameters (green – KPZ scaling, yellow – KPZ scaling with holes and detachments, red – jammed, collective movement; brown – jammed, no movement.

## Supplementary Movie Legends

**Supplementary Movie 1:** Invasion of A431 human skin carcinoma cells in ordered vs. disordered environments. Cells express cytoplasmic mCherry (blue) and nuclear H2B-GFP (green).

**Supplementary Movie 2:** Invasion of A431 human skin carcinoma cells in free environments. Cells express cytoplasmic mCherry (blue) and nuclear H2B-GFP (green).

**Supplementary Movie 3:** Invasion of EPCad^-/-^ A431 human skin carcinoma cells in ordered vs. disordered environments. Cells express cytoplasmic mCherry (blue) and nuclear H2B-GFP (green).

**Supplementary Movie 4:** Simulations of cells with different fluidity number (i.e. with varying cell-cell interaction strength) in homogeneous environment showing different level of cell rearrangements. Cells are represented as grey beads.

**Supplementary Movie 5:** Simulations of cells migrating in different environments (homogeneous, ordered and disordered. Cells are represented as grey and pillars as black beads.

**Supplementary Movie 6:** Simulations of EPCad^-/-^ cells (lower cell-cell adhesion) invading different types of environments. Simulations of cells migrating in different environments (homogeneous, ordered and disordered. Cells are represented as grey and pillars as black beads.

**Supplementary Movie 7:** Simulations of cell invasion in spatially structured environments which do not increase detachment rate (regular with smaller pore, sinusoid, stepwise). Cells are represented as grey and pillars as black beads.

**Supplementary Movie 8:** Simulation and experiment of cell invasion in sinusoid perpendicular pattern, which creates localized interface protrusions and detachments. Cells in experiment express cytoplasmic mCherry (blue) and nuclear H2B-GFP (green). Cells in simulations are represented as grey and pillars as black beads.

## Notes

### Competing Interest Statement

The authors have declared no competing interest.

### Summary of Updates

New supplementary figure panels to look at the scaling exponents of front invasion. rewriting of the main text

